# Comparison of directional random walk and weighted least squares modeling of sparse fossil data

**DOI:** 10.64898/2026.06.26.734751

**Authors:** Rolf Ergon

## Abstract

The general random walk model (GRW) of Hunt (2006) is used to infer directional evolution in mean trait values from sparse fossil data by modeling phenotypic change as the accumulated result of small steps with mean step sizes and step variances. Using simulations and real data cases, Ergon (2026) showed that the step variances can be estimated reasonably well only when the mean trait values have small measurement errors, while for fossil data with realistic measurement errors they appear to be extremely difficult to find, and they are often found to be negative. In the simulations Ergon (2026) assumed that the true phenotypic mean values were known. Here, I essentially repeat these simulations under the assumption that only mean trait values with large measurement errors are known, and based on weighted mean squared error (WMSE) comparisons the conclusion is that weighted least squares (WLS) is a better method than GRW. A second conclusion is that WLS is a better method also in the possibly rare cases with large measurement errors where the GRW parameters are estimated well. The GRW method is simply not flexible enough to handle such cases. A third conclusion is that Akaike Information Criterion (AIC) results for GRW models with large measurement errors relative to the step variance may be overly optimistic.

## 1. Introduction

Hunt (2006) developed what he called a general random walk (GRW) model for analyses of fossil time series, later referred to as a model for directional evolution (Hunt, 2012). This model assumes a random walk process where at each timestep an increment of evolutionary change in a phenotypic mean trait value is drawn at random from a distribution of evolutionary steps, and that this incremental change is normally distributed with mean value *b*_*GRW*_ and variance 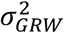. For an observed evolutionary mean trait change Δ*Y* over T time steps the log-likelihood function for a GRW process is given by (Hunt, 2006)

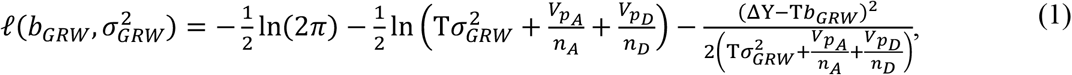

where *n*_*A*_ and *n*_*D*_ are the numbers of observed ancestors and descendants, respectively, while *V*_*pA*_ and *V*_*pD*_ are the corresponding population phenotypic variances. Note that Hunt (2006) used the notation *μ*_*step*_ for the mean step size, and that the directional slope is *b*_*GRW*_ = *cμ*_*step*_, where *c* can be chosen freely to for example *c* = 1. With *N* irregular and sparse samples of mean trait values, multiple ancestor-descendant mean trait differences over *N* − 1 evolutionary transitions may be used jointly to estimate *b*_*GRW*_ and 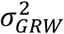by summing the log-likelihoods according to Eq. (1) over the transitions, i.e., by use of

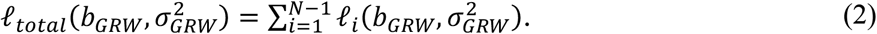

The step parameters in Eq. (2) can be found by maximizing 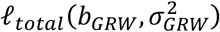, i.e., by maximum likelihood estimation, and the estimated step size 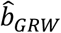 will then be a measure of directional change over time.

For a single transition it is obvious from Eq. (1) that 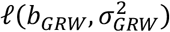 is maximized when 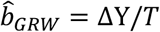, and as shown in simulations in Section 3, *b*_*GRW*_ can be fairly well estimated also from data over several transitions. From Eq. (1) it is also obvious that it is difficult to estimate 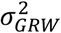 in cases where 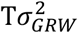 is small compared to *V*_*p*_⁄*n* for the samples, i.e., when the measurement errors are large. This problem was studied in Ergon (2026), assuming that the true mean trait values were known, and that resulted in two major conclusions. First, in many cases with realistic measurement errors the estimated values of 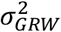 become negative, such that one must set 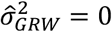. In such cases the estimated response slope 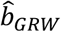 will be non-optimal as compared to generalized least squares (GLS) or weighted least squares (WLS) estimates. In Ergon (2026) I described four such cases with real data. Second, also in cases with 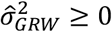, large measurement errors will lead to non-optimal 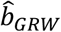estimates.

Hunt (2006) tested the GRW model with what appears to be unrealistically small phenotypic variances *V*_*pA*_ = *V*_*pD*_ = 1 and *n*_*A*_ = *n*_*D*_ = 30, and he noted that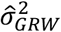 in some cases could be negative. In Ergon (2026) I essentially repeated Hunt’s simulations, but with *V*_*pA*_ = *V*_*pD*_ = 400, and found 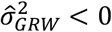 in around 50% of the realizations.

In Ergon (2026) I thus studied the effects of poor estimates of 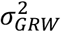, which is an obvious practical problem since 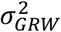 in Eq. 1 only appears in the sum 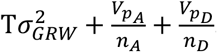. In the present paper I essentially repeat the simulations in Ergon (2026), but with use of measured instead of true mean trait values, and the conclusions are the same. I also ask whether good estimates of *b*_*GRW*_ and 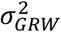 would guarantee good estimates of the directional slope, and as shown in simulations the answer is no. The GRW model is simply not flexible enough to handle large measurement errors, such that also in realizations where the GRW parameter estimates happen to be good, the optimal results are still found by use of WLS. The conclusions are reached by comparisons using weighted mean squared errors (WMSE) values, but results by use of the Akaike Information Criterion (AIC) is also discussed. I found, however, that AIC results for GRW models with large measurement variances as compared to 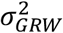 may be too optimistic, possibly owing to the dependence between adjacent trait differences because they share a sample mean and its sampling error (Hunt, 2006).

## 2. The Akaike Information Criterion for GRW and WLS

As explained in more detail in Hunt (2006) and Ergon (2026), the Akaike Information Criterion with short data corrections for a GRW model is found as

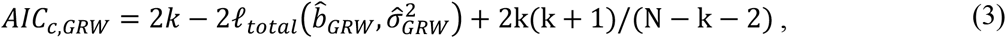

Where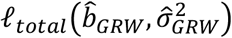is the log-likelihood function over all transitions, and where k = 2 is the number of estimated parameters, while *N* − 1 = 9 is the number of transitions.

For computation of *AIC*_*c,WLS*_, Banks and Joyner (2017) assumed a statistical model for a random variable

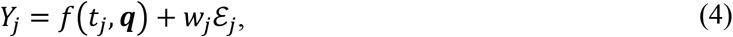

where *w*_*j*_ are known weights and where ℰ_*j*_ for *j* = 1, 2, …, *N* is i.i.d. N(0, σ^2^), while ***q*** is a parameter vector. Under this model they found

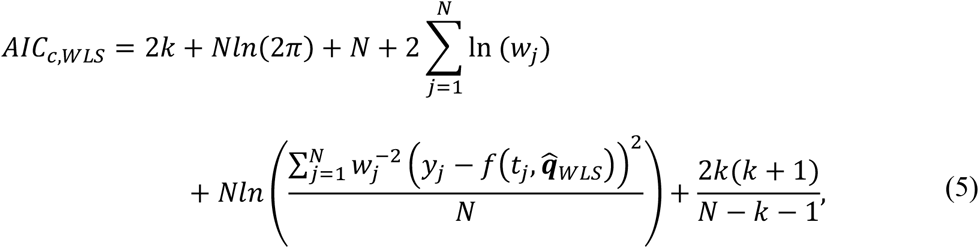

where using the notation in Ergon (2026) 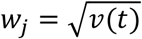 and 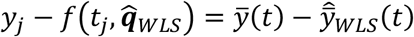, while since also σ^2^ must be estimated *k* = 3.

## 3. Old and new simulations

In Ergon (2026) I generated true responses as

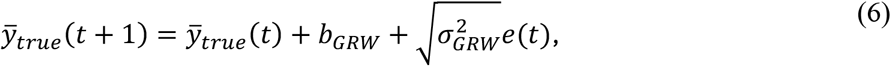

with 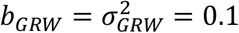, and with *e*(*t*) as a normally distributed random numbers with variance one. I then added measurement errors to create ten samples 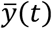. For computation of WMSE values I assumed 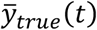 to be known.

Here, I extend the simulations under the assumption that only 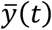 is known, i.e., with measurement errors, although I compute a reference prediction slope 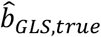 using 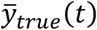 as described in Ergon (2026). For model comparisons I primarily use WMSE values, although also *AIC*_*c*_ results are discussed. I use both regular sampling and irregular sampling with variable numbers of samples behind each mean value as described in Ergon (2026).

For GRW the maximum likelihood search for step variance is limited to 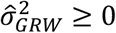. From the estimated prediction slope 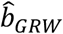 follows

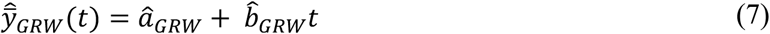

by fitting a parameter *a*_*GWR*_ to the data by a weighted least squares approach, making use of the diagonal matrix ***V*** with elements *v*_*t*_ = *V*_*p*_⁄*n*_*t*_. From 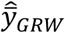 follows

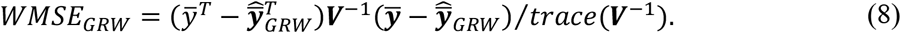

For WLS the parameter estimates are found as

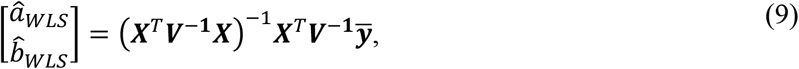

where 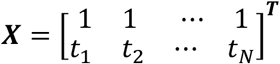 and 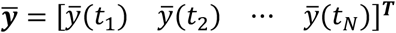, and from this follows

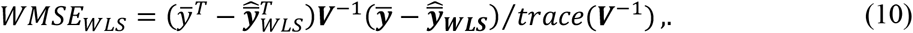

### Remark 1

One could argue that the diagonal matrix ***V*** in Eq. (8) should be replaced by ***C*** + ***V***, where ***C*** is the covariance matrix as found in Appendix A, but that would make fair comparisons between *WMSE*_*GRW*_ and *WMSE*_*WLS*_ impossible because of different normalizations.

### Remark 2

It should be noted that *AIC*_*c,WLS*_ according to Eq. (5) is not affected by various forms of normalization in the computation of *WMSE*_*WLS*_. Any normalization will be compensated by the estimated value of σ^2^, such that questions of normalization are irrelevant.

## 4. Simulation results

### 4.1 Prediction slope and WMSE results

Prediction slopes 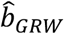 are computed by maximizing 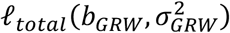as given in Eq. (2) under the constraint 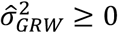 while 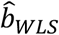 is computed according to Eq. (9), and the results are shown in Table 1. Fig. 1 shows histograms over prediction errors in 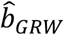 and 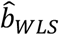 as found by comparisons with predictions based on true 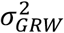 and 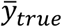 values, where 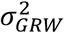enters the computations via the covariance matrix as described in Appendix A. As can be seen in Fig. 1, the prediction errors for 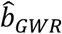 with 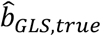 as reference are large compared to the corresponding errors for 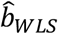. As can be seen in Table 1, all realizations resulted in *WMSE*_*WLS*_ < *WMSE*_*GRW*_.

**Table 1.** Simulation results given as *mean* ± *SE* from *1,000* realizations.

| | $V_p = 1$<br>regular sampling | $V_p = 400$<br>regular sampling | $V_p = 400$<br>irregular sampling |
| --- | --- | --- | --- |
| $\hat{\sigma}_{step}^2$ | $0.0900 \pm 0.0467$ | $0.1144 \pm 0.1528$ | $0.1210 \pm 0.2175$ |
| $\hat{\mu}_{step} = \hat{b}_{GRW}$ | $0.1001 \pm 0.0102$ | $0.1000 \pm 0.0121$ | $0.1002 \pm 0.0150$ |
| $\hat{b}_{WLS}$ | $0.1001 \pm 0.0108$ | $0.1001 \pm 0.0120$ | $0.0998 \pm 0.0119$ |
| $WMSE_{GWR}$ | $7.4 \pm 4.9$ | $19.1 \pm 10.1$ | $25.5 \pm 18.2$ |
| $WMSE_{WLS}$ | $6.5 \pm 4.3$ | $16.8 \pm 8.6$ | $16.6 \pm 9.4$ |
| $WMSE_{WLS} < WMSE_{GRW}$ | 100% | 100% | 100% |

**Figure 1.**
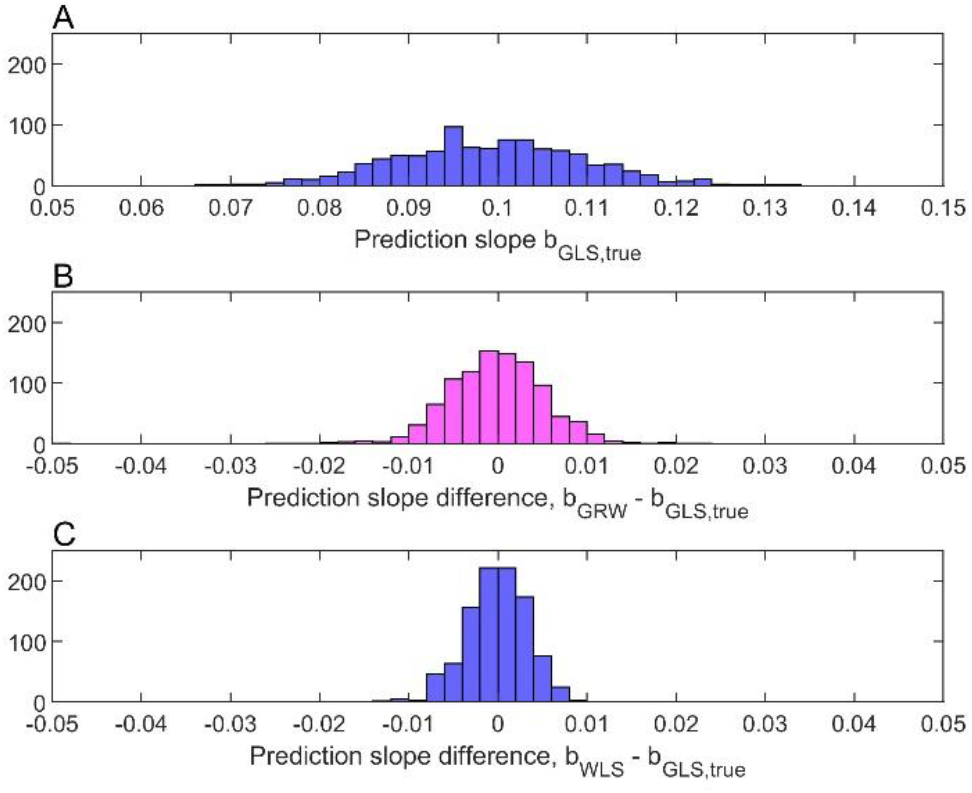
Histogram for 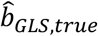 using *V*_*p*_ = 400 under the assumption that 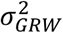 and 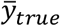 are known (panel A), and histograms for prediction errors 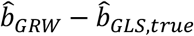 (panel B) and 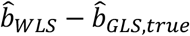 (panel C) based on 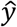 with measurement errors.

### 4.2 Results with good parameter estimates

The results in Table 1 raises the question of results for the few realizations with *V*_*p*_ = 400 where 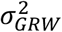 is estimated well, and the answer is given in Table 2. Here, results for small errors in 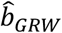 and for small errors in both 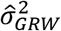and 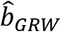 are included. The results are essentially the same as in Table 1, and there is thus no help in having good parameter estimates. The GRW model is just not flexible enough to handle large measurement errors.

**Table 2.**
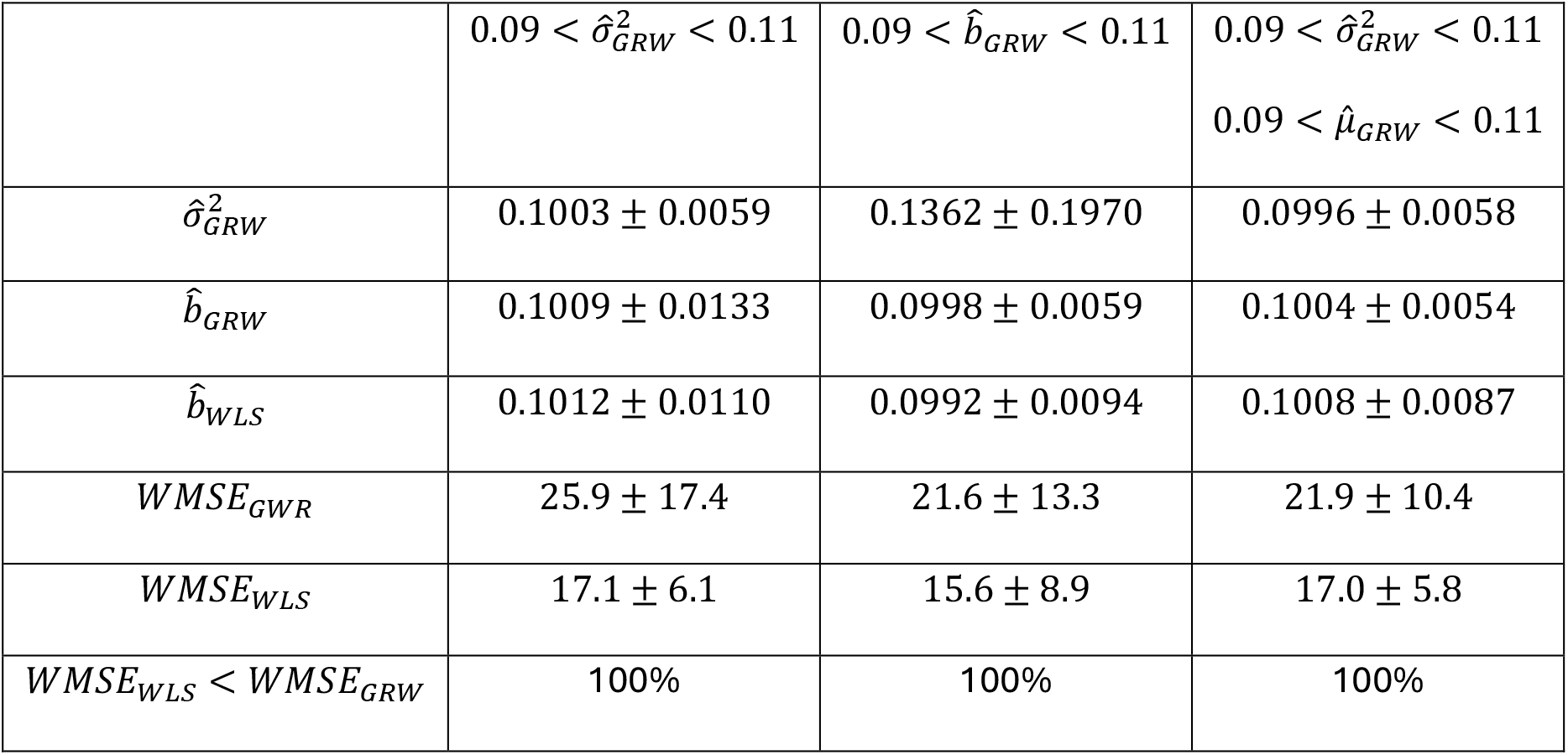
Simulation results given as *mean* ± *SE* from 100 realizations with *V*_*p*_ = 400 and small errors in 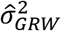 and 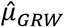, separately, and small errors in both 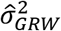 and 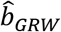.

| | $0.09 < \hat{\sigma}_{GRW}^2 < 0.11$ | $0.09 < \hat{b}_{GRW} < 0.11$ | $0.09 < \hat{\sigma}_{GRW}^2 < 0.11$<br>$0.09 < \hat{\mu}_{GRW} < 0.11$ |
| --- | --- | --- | --- |
| $\hat{\sigma}_{GRW}^2$ | $0.1003 \pm 0.0059$ | $0.1362 \pm 0.1970$ | $0.0996 \pm 0.0058$ |
| $\hat{b}_{GRW}$ | $0.1009 \pm 0.0133$ | $0.0998 \pm 0.0059$ | $0.1004 \pm 0.0054$ |
| $\hat{b}_{WLS}$ | $0.1012 \pm 0.0110$ | $0.0992 \pm 0.0094$ | $0.1008 \pm 0.0087$ |
| $WMSE_{GWR}$ | $25.9 \pm 17.4$ | $21.6 \pm 13.3$ | $21.9 \pm 10.4$ |
| $WMSE_{WLS}$ | $17.1 \pm 6.1$ | $15.6 \pm 8.9$ | $17.0 \pm 5.8$ |
| $WMSE_{WLS} < WMSE_{GRW}$ | 100% | 100% | 100% |

### 4.3 AIC comparisons

#### 4.3.1 Simulation results

Ideally, the GRW and WLS models should be compared by use of AIC. This is not meaningful, however, since as shown in Table 3 large ratios 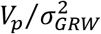appear to affect *AIC*_*c,GRW*_ according to Eq. (3). This may possibly be caused by the “dependence between adjacent trait differences because they share a sample mean and its sampling error”, as discussed in Hunt (2006).

**Table 3.** Examples of *AIC*_*c*_ and WMSE differences for realizations with *V*_*p*_ = 400 and irregular sampling.

| $\hat{\sigma}_{GRW}^2$ | $AIC_{c,GRW} - AIC_{c,WLS}$ | $WMSE_{GRW} - WMSE_{WLS}$ |
| --- | --- | --- |
| 0.0000 | -4.0176 | 12.6682 |
| 0.7841 | 1.6268 | 39.7548 |
| 0.0000 | 0.4674 | 0.0532 |
| 0.1246 | -3.8024 | 3.1632 |
| 0.0401 | -0.6048 | 0.5177 |
| 0.0000 | -6.0595 | 9.3068 |
| 0.3137 | -6.9934 | 33.1775 |
| 0.0000 | 2.6067 | 0.0022 |
| 0.5319 | -3.6179 | 6.3324 |
| 0.0000 | 1.4386 | 16.3187 |
| 0.0154 | -1.2282 | 0.1281 |

In Table 3, *AIC*_*c,GRW*_ < *AIC*_*c,WLS*_ for some realizations where 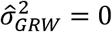, which should not be possible since *AIC*_*c,WLS*_ then follows from the best linear unbiased estimate (BLUE) of *b*_*WLS*_.

GRW with 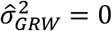is a purely deterministic model, and it cannot then be better than the optimal WLS model. In one example realization with 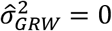(red numbers) the results are *AIC*_*c,GRW*_ − *AIC*_*c,WLS*_ ≈ −6 and *WMSE*_*GRW*_ − *WMSE*_*WLS*_ ≈ 9, and such a large discrepancy between AIC and WMSE model quality results cannot be explained by differences in short data corrections. For this phenomenon to occur with 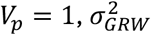 must be reduced to 0.001, which might explain why Hunt (2006) found the problem unimportant. Note that values with 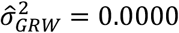in Table 3 imply that 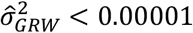, and such values close to the lower search limit zero were found in around 45% of the realizations. Also note that all realizations in Table 3 have *WMSE*_*GRW*_ > *WMSE*_*WLS*_.

Although *AIC*_*c,GRW*_ < *AIC*_*c,WLS*_ for some realizations with 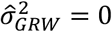, that is not always the case. In another example realization in Table 3 (orange numbers) we find *AIC*_*c,GRW*_ − *AIC*_*c,WLS*_ ≈ 2.6 and *WMSE*_*GRW*_ − *WMSE*_*WLS*_ ≈ 0.

#### 4.3.2 Real data results

##### Ergon (2026) found prediction slope and WMSE results for four real data cases, all with

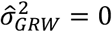. These results are summarized in Table 4, where also AIC results are included. Here, *AIC*_*c,GWR*_ is found according to Eq. (3), while *AIC*_*c,GWR,WLS*_ is found by applying Eq. (5) on the *WMSE*_*GWR*_ results. In the Bryozoan and Stickleback cases *WMSE*_*GRW*_ ≈ *WMSE*_*WLS*_ and *AIC*_*c,GRW,WLS*_ ≈ *AIC*_*c,WLS*_, indicating that the GRW and WLS models are equally good. According to the *AIC*_*c,GRW*_ results, however, GRW is a much poorer model than WLS, which again shows that Eq. (3) cannot be trusted.

**Table 4.** Real data results with 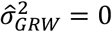 given as *mean* ± *SE* from *1,000* realizations.

|  | Bryozoan | Ostracod 1 | Ostracod 2 | Stickleback |
| --- | --- | --- | --- | --- |
| $\hat{b}_{GRW}$ | 0.1441 | 0.0312 | 0.0254 | 1.2531 |
| $\hat{b}_{WLS}$ | 0.1519 | 0.0402 | 0.0434 | 1.2605 |
| $WMSE_{GRW}$ | 0.00192 | 0.00033 | 0.00109 | 0.00040 |
| $WMSE_{WLS}$ | 0.00190 | 0.00024 | 0.00061 | 0.00040 |
| $AIC_{c,GRW}$ | -10.2 | -22.9 | -4.2 | -104 |
| $AIC_{c,GRW,WLS}$ | -17.5 | -39.6 | -21.9 | -121 |
| $AIC_{c,WLS}$ | -17.6 | -42.6 | -27.8 | -121 |

For completeness, Table 5 shows results when 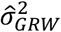 is allowed to be negative. In the Bryozoan case GRW has the lowest *AIC*_*c*_ result, which appears to be plausible since 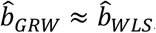, but a negative variance is of course not possible. Also note the large increase in 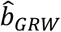 in the Stickleback case.

**Table 5.**
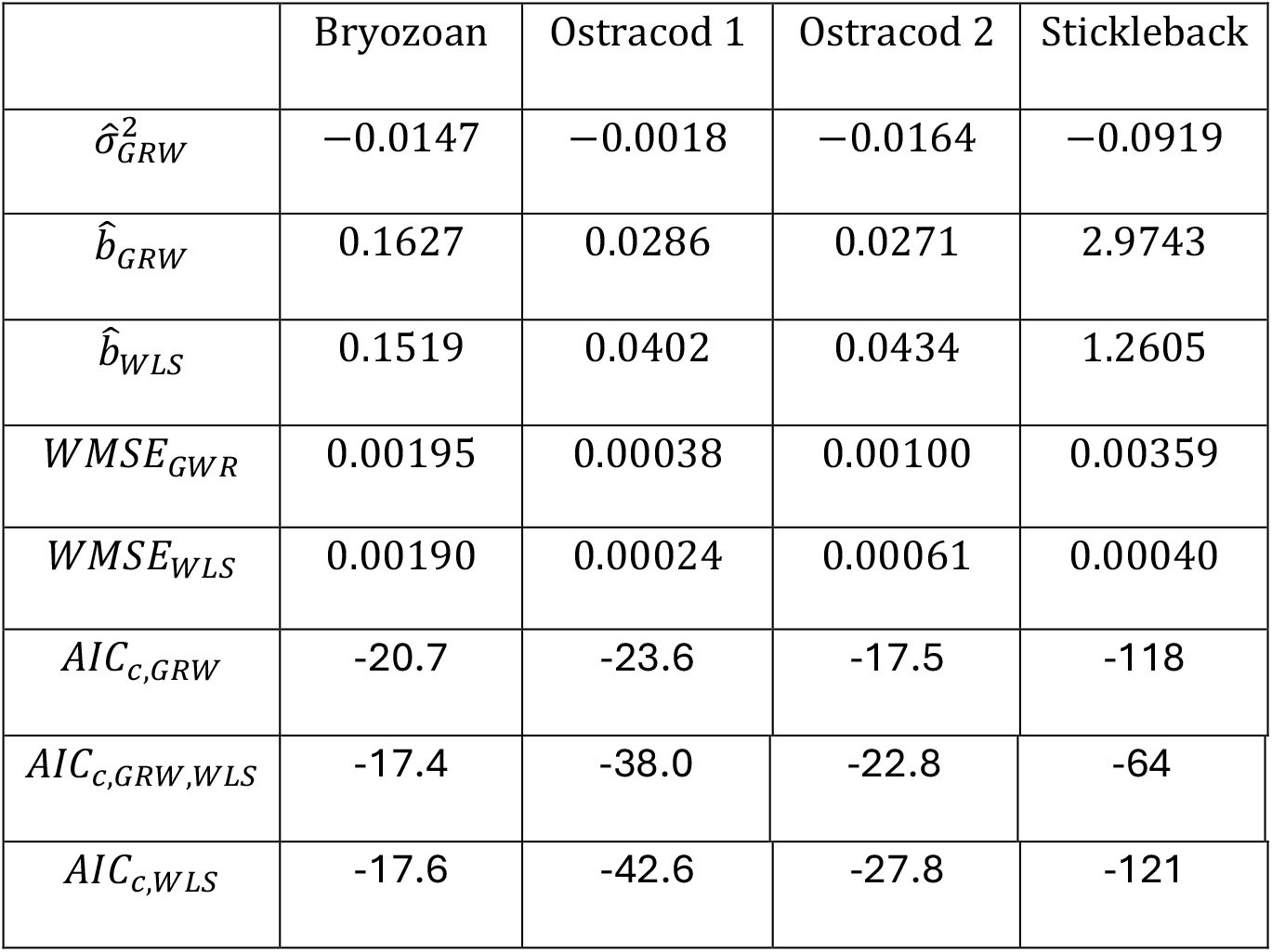
Real data results with 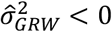 given as *mean* ± *SE* from *1,000* realizations.

## 5. Summary and discussion

In real data cases it is obviously possible to compute and compare *WMSE*_*GRW*_ and *WMSE*_*WLS*_, and the intention of the simulations is thus to indicate what may be expected under the assumptions of an underlying GRW model and large measurement errors. Ideally, the models should be compared by use of AIC, but as found in Section 4 and discussed below this appears not to be meaningful when the measurement variances are large compared to the incremental step variance 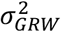.

There are three main conclusions to be drawn from the simulation results. First, and as shown in Table 1, WLS should be expected to be the best model in all practical cases with large measurement errors compared to 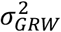in an underlying GRW model. Second, as shown in Table 2, it does not help if the estimates of 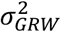 and/or *b*_*GRW*_ should happen to be good. The GRW model is just not flexible enough to handle large measurement errors. Third, as shown in Tables 3 and 4, *AIC*_*c,GRW*_ values are not reliable when the measurement variances are large compared to the incremental step variance, see Subsection 4.3 for a more detailed discussion.

It is a serious problem that *AIC*_*c,GRW*_ values with large measurement variances relative to 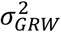in some cases appear to be erroneous, and further research of this topic is thus important. One aspect of the problem is that correlated terms in 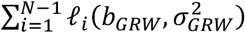 require a modification of the resulting 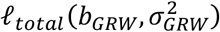 in Eq. (2).

## Supporting information

MATAB Code

## Appendix A. Generalized least squares (GLS) results

Assuming an underlying GRW model, with estimated parameter values 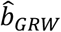and 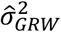, *WMSE*_*GRW*_ and *WMSE*_*GLS*_ can be found from Eqs. (8) and (10) by replacing ***V***^−1^ with **(*C***+***V***)^−1^, where (Ergon, 2026),

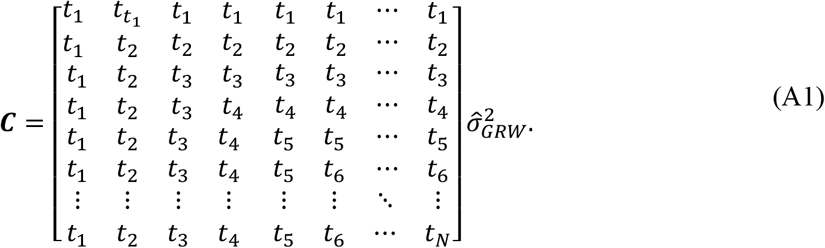

Results corresponding to Table 1 are shown in Table A1. Especially note that the mean values for both *WMSE*_*GWR*_ and *WMSE*_*GLS*_ for *V*_*p*_ = 1 are reduced, but at the cost of increased SE values. Also note that *WMSE*_*GLS*_ > *WMSE*_*GRW*_for a few realizations.

**Table A1.**
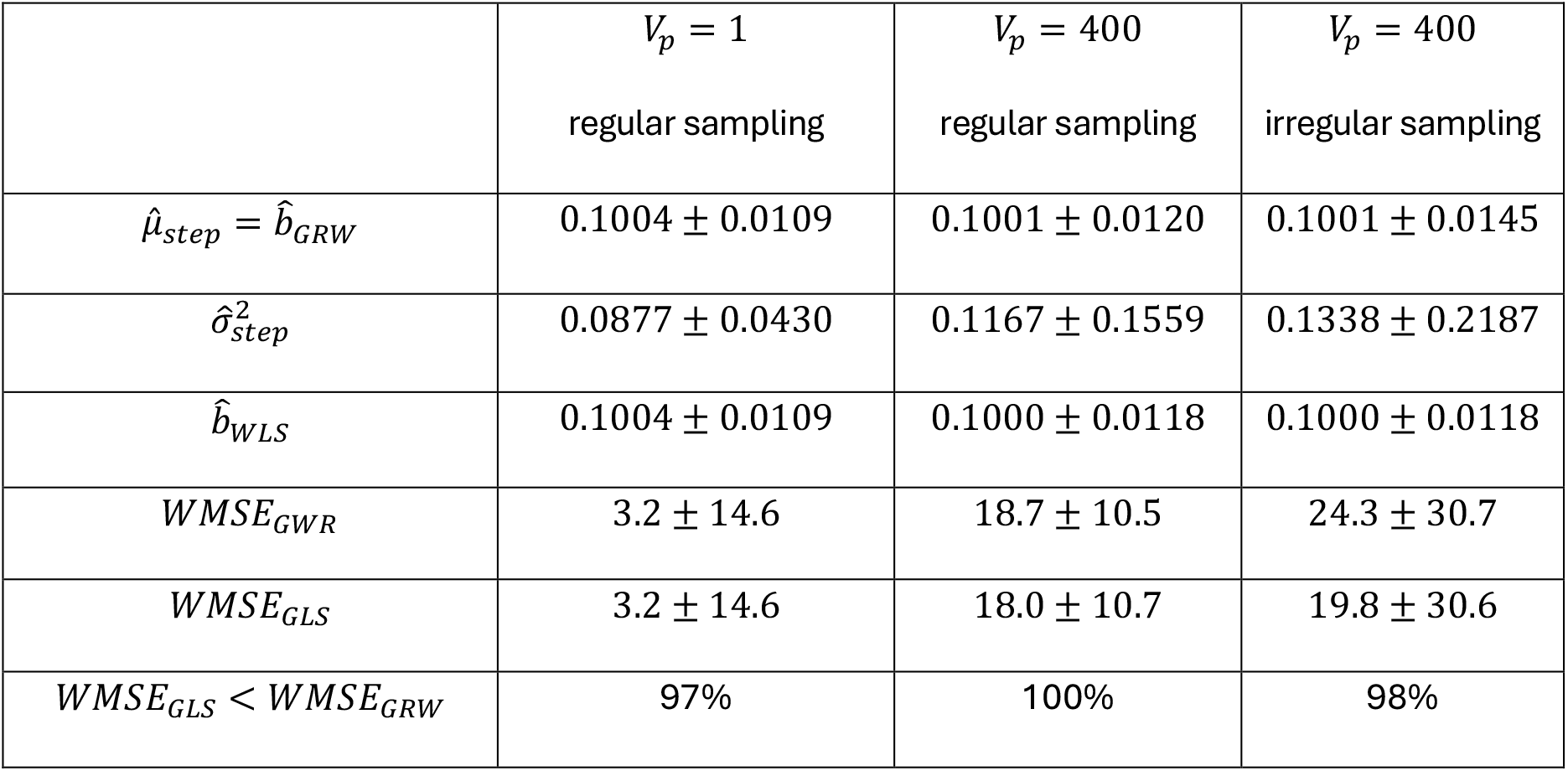
Simulation results given as *mean* ± *SE* from 1,000 realizations.

## Acknowledgment

I thank University of South-Eastern Norway for support and funding.

## References

Banks, H.T., and Joyner, M.L. (2017). AIC under the framework of least squares estimation. Applied Mathematics Letters. 10.1016/j.aml.2017.05.005

Ergon, R. (2026). A critical look at directional random walk modeling of sparse fossil data. Ecology and Evolution, 2026; 16:e73669 10.1002/ece3.73669

Hunt, G. (2012). Measuring rates of phenotypic evolution and the inseparability of tempo and Mode. Paleobiology, 38(3), 2012, pp. 351–373. paleo_Hunt_2012_Pb_rates.pdf

Hunt, G. (2006). Fitting and comparing models of phyletic evolution: random walks and beyond. Paleobiology 32: 578–601.

