## Supplementary material for "Comparison of directional random walk and weighted least squares modeling of sparse fossil data": MATAB Code

### MATLAB Code

### for

Rolf Ergon

University of South-Eastern Norway

June 25, 2026

#### Figure 1

```
clear


mu_step=0.1;
var_step=0.1;
% var_step=0;
Vp=100;
count_negvar=0;
count_WLSbest=0;
y0=0;
M=1000
for m=1:M
%% Generate data
y=y0*zeros(1,1000);
for t=2:1000
   tplot(t)=t;
   y(t)=y(t-1)+mu_step+sqrt(var_step)*randn;
end
tlabel=zeros(1,10);
ylabell=zeros(1,10);
err=zeros(1,10);


j=1;
for j=1:10
```

```
    for t=1:1000
        if t/100==j
            tlabel(j)=t-99*rand;
            j=j+1;
        end
    end
end
tlabel=round(tlabel);
for j=1:10
    n(1,j)=57*rand;
end
n=round(n)+3*ones(1,10);

j=1;
for t=1:1000
    if t==tlabel(j)
        individuals=sqrt(Vp)*randn(n(j),1);
        Mean(1,j)=mean(individuals);
        ytrue(j)=y(t);
        ylabell(j)=y(t)+Mean(1,j);
        err(j)=sqrt(Vp/n(j));
        if j==10 break
        end
        j=j+1;
    end
end

v=err.^2;
V=diag(v);

%% Covariance for GLS model
tv=tlabel;
Cov=[tv(1) tv(1) tv(1) tv(1) tv(1) tv(1) tv(1) tv(1) tv(1) tv(1)
     tv(1) tv(2) tv(2) tv(2) tv(2) tv(2) tv(2) tv(2) tv(2) tv(2)
     tv(1) tv(2) tv(3) tv(3) tv(3) tv(3) tv(3) tv(3) tv(3) tv(3)
     tv(1) tv(2) tv(3) tv(4) tv(4) tv(4) tv(4) tv(4) tv(4) tv(4)
     tv(1) tv(2) tv(3) tv(4) tv(5) tv(5) tv(5) tv(5) tv(5) tv(5)
     tv(1) tv(2) tv(3) tv(4) tv(4) tv(5) tv(6) tv(6) tv(6) tv(6)
```

```
    tv(1) tv(2) tv(3) tv(4) tv(5) tv(6) tv(7) tv(7) tv(7) tv(7)
    tv(1) tv(2) tv(3) tv(4) tv(5) tv(6) tv(7) tv(8) tv(8) tv(8)
    tv(1) tv(2) tv(3) tv(4) tv(5) tv(6) tv(7) tv(8) tv(9) tv(9)
    tv(1) tv(2) tv(3) tv(4) tv(5) tv(6) tv(7) tv(8) tv(9) tv(10)];

%% GRW

n_samples=ones(1,10);
var_samples=err.^2;

c=1;
Tsample=c*tlabel;

for i=1:9
   dT(i)=-Tsample(i)+Tsample(i+1);
   dX(i)=ylabell(i+1)-ylabell(i);
   nA(i)=n_samples(i);
   nD(i)=n_samples(i+1);
   varA(i)=var_samples(i);
   varD(i)=var_samples(i+1);
end

% Contraints
mustep_min=-100; mustep_max=100;
varstep_min=0; varstep_max=100;
par_lb=[mustep_min varstep_min];
par_ub=[mustep_max varstep_max];

% fmincon
par_guess=[0 0];
Aineq=[]; Bineq=[]; Aeq=[]; Beq=[];
fun_objective_handle=...
   @(par)fun_objective(par,dT,dX,varA,nA,varD,nD);
[par_opt,fval,exitflag,output,lambda,grad,hessian] =...
fmincon(fun_objective_handle,par_guess,Aineq,Bineq,Aeq,Beq,par_lb,par_ub);

mustep(m)=par_opt(1);
varstep(m)=par_opt(2);
```

```
if varstep(m)<0
   varstep(m)=0;
end

b_GRW(m)=mustep(m);

for i=1:9
   varterm(i)=dT(i)*varstep(m)+varA(i)/nA(i)+varD(i)/nD(i);
   Li(i)=-log(2*pi)/2-log(varterm(i))/2-(dX(i)-dT(i)*mustep(m))^2/(2*varterm(i));
end

L=sum(Li);

%% GLS based on var_step and ytrue

Ctrue=Cov*var_step;
CVtrue=inv(Ctrue+V);

X=[ones(10,1) tlabel'];
bls=inv(X'*CVtrue*X)*X'*CVtrue*ytrue';

a_GLS=bls(1);
b_GLS(m)=bls(2);
for i=1:10
   yhat_GLS(i)=a_GLS+b_GLS(m)*tlabel(i);
end

b_GLS_true(m)=b_GLS(m);

if varstep(m)==0
   b_GLS_true(m)=NaN;
end

%% WLS

X=[ones(10,1) tlabel'];
bls=inv(X'*inv(V)*X)*X'*inv(V)*ylabell';
```

```
a_WLS=bls(1);
b_WLS(m)=bls(2);
for i=1:10
   yhat_WLS(i)=a_WLS+b_WLS(m)*tlabel(i);
end

end

figure(1)
subplot(3,1,1)
histogram(b_GLS_true,80,'BinWidth',0.002,'FaceColor','b')
ax = findobj(subplot(3,1,1),'Type','Axes');
for i = 1:length(ax)
   ylim(ax(i),[0 250]);
   xlim(ax(i),[0.05 0.15]);
end
xlabel('Prediction slope b_G_L_S_,_t_r_u_e')
text(0.05,280,'A','FontSize',12)

subplot(3,1,2)
histogram(b_GRW-b_GLS_true,80,'BinWidth',0.002,'FaceColor','m')
ax = findobj(subplot(3,1,2),'Type','Axes');
for i = 1:length(ax)
   ylim(ax(i),[0 250]);
   xlim(ax(i),[-0.05 0.05]);
end
xlabel('Prediction slope difference, b_G_R_W - b_G_L_S_,_t_r_u_e')
text(-0.05,280,'B','FontSize',12)

subplot(3,1,3)
histogram(b_WLS-b_GLS_true,80,'BinWidth',0.002,'FaceColor','b')
ax = findobj(subplot(3,1,3),'Type','Axes');
for i = 1:length(ax)
   ylim(ax(i),[0 250]);
   xlim(ax(i),[-0.05 0.05]);
end
xlabel('Prediction slope difference, b_W_L_S - b_G_L_S_,_t_r_u_e')
text(-0.05,280,'C','FontSize',12)
```

```
%% Parameter search

function f = fun_objective(par,dT,dX,varA,nA,varD,nD)

mustep=par(1);

varstep=par(2);


for i=1:9

   varterm(i)=dT(i)*varstep+varA(i)/nA(i)+varD(i)/nD(i);

   Li(i)=-log(2*pi)/2-log(varterm(i))/2-(dX(i)-dT(i)*mustep)^2/(2*varterm(i));

end


L=sum(Li);


f=-L;


end
```

#### Table 1

```
clear


mu_step=0.1;

var_step=0.1;

Vp=400;

count=0;

y0=0;

M=1000

for m=1:M

%% Generate data

y=y0*zeros(1,1000);

for t=2:1000

   tplot(t)=t;

   y(t)=y(t-1)+mu_step+sqrt(var_step)*randn;

end

tlabel=zeros(1,10);

ylabell=zeros(1,10);

err=zeros(1,10);
```

```
% j=1;                % Use for regular sampling
% for t=1:1000
%    if t/100==j
%       tlabel(j)=t-50;
%       j=j+1;
%    end
% end
% tlabel=round(tlabel);
% n=30*ones(1,10);

j=1;              % Use for irregular sampling
for j=1:10
   for t=1:1000
      if t/100==j
         tlabel(j)=t-99*rand;
         j=j+1;
      end
   end
end
tlabel=round(tlabel);
for j=1:10
   n(1,j)=57*rand;
end
n=round(n)+3*ones(1,10);

j=1;
for t=1:1000
   if t==tlabel(j)
      individuals=sqrt(Vp)*randn(n(j),1);
      Mean(1,j)=mean(individuals);
      ytrue(j)=y(t);
      ylabell(j)=y(t)+Mean(1,j);
      err(j)=sqrt(Vp/n(j));
      if j==10 break
      end
      j=j+1;
   end
```

```
end

v=err.^2;
V=diag(v);

%% Covariance matrix for use in Appendix A
tv=tlabel;
Cov0=[tv(1) tv(1) tv(1) tv(1) tv(1) tv(1) tv(1) tv(1) tv(1) tv(1)
    tv(1) tv(2) tv(2) tv(2) tv(2) tv(2) tv(2) tv(2) tv(2) tv(2)
    tv(1) tv(2) tv(3) tv(3) tv(3) tv(3) tv(3) tv(3) tv(3) tv(3)
    tv(1) tv(2) tv(3) tv(4) tv(4) tv(4) tv(4) tv(4) tv(4) tv(4)
    tv(1) tv(2) tv(3) tv(4) tv(5) tv(5) tv(5) tv(5) tv(5) tv(5)
    tv(1) tv(2) tv(3) tv(4) tv(4) tv(5) tv(6) tv(6) tv(6) tv(6)
    tv(1) tv(2) tv(3) tv(4) tv(5) tv(6) tv(7) tv(7) tv(7) tv(7)
    tv(1) tv(2) tv(3) tv(4) tv(5) tv(6) tv(7) tv(8) tv(8) tv(8)
    tv(1) tv(2) tv(3) tv(4) tv(5) tv(6) tv(7) tv(8) tv(9) tv(9)
    tv(1) tv(2) tv(3) tv(4) tv(5) tv(6) tv(7) tv(8) tv(9) tv(10)];

%% GRW

n_samples=ones(1,10);
var_samples=err.^2;

c=1;
Tsample=c*tlabel;

for i=1:9
   dT(i)=-Tsample(i)+Tsample(i+1);
   dX(i)=ylabell(i+1)-ylabell(i);
   nA(i)=n_samples(i);
   nD(i)=n_samples(i+1);
   varA(i)=var_samples(i);
   varD(i)=var_samples(i+1);
end

% Contraints
mustep_min=-100; mustep_max=100;
varstep_min=0; varstep_max=100;
```

```
par_lb=[mustep_min varstep_min];
par_ub=[mustep_max varstep_max];

% fmincon
par_guess=[0 0];
Aineq=[]; Bineq=[]; Aeq=[]; Beq=[];
fun_objective_handle=...
   @(par)fun_objective(par,dT,dX,varA,nA,varD,nD);
[par_opt,fval,exitflag,output,lambda,grad,hessian] =...
fmincon(fun_objective_handle,par_guess,Aineq,Bineq,Aeq,Beq,par_lb,par_ub);

mustep(m)=par_opt(1);
varstep(m)=par_opt(2);

varstepestimated(m)=varstep(m);

if varstep(m)<0
   varstep(m)=0;
end

% Cov=Cov0*varstep(m);    % For use in Appendix A
% V=Cov+V;

X=ones(10,1);
b_GRW(m)=mustep(m);
a_GRW(m)=inv(X'*inv(V)*X)*X'*inv(V)*(ylabell'-b_GRW(m)*tlabel');

for j=1:10
   yhat_GRW(j)=a_GRW(m)+b_GRW(m)*tlabel(j);
end

WMSE_GRW(m)=(ylabell-yhat_GRW)*inv(V)*(ylabell-yhat_GRW)'/trace(inv(V));

%% WLS

X=[ones(10,1) tlabel'];
bls=inv(X'*inv(V)*X)*X'*inv(V)*ylabell';
a_WLS(m)=bls(1);
```

```
b_WLS(m)=bls(2);
for i=1:10
   yhat_WLS(i)=a_WLS(m)+b_WLS(m)*tlabel(i);
end

WMSE_WLS(m)=(ylabell-yhat_WLS)*inv(V)*(ylabell-yhat_WLS)'/trace(inv(V));

if WMSE_WLS(m)<WMSE_GRW(m)
   count=count+1;
end

end

Mstep=[mean(mustep) std(mustep)]
Varstep=[mean(varstep) std(varstep)]
B1=[mean(b_WLS) std(b_WLS)]
WMSEGRW=[mean(WMSE_GRW) std(WMSE_GRW)]
WMSEWLS=[mean(WMSE_WLS) std(WMSE_WLS)]

100*count/M

%% Parameter search
function f = fun_objective(par,dT,dX,varA,nA,varD,nD)
mustep=par(1);
varstep=par(2);

for i=1:9
   varterm(i)=dT(i)*varstep+varA(i)/nA(i)+varD(i)/nD(i);
   Li(i)=-log(2*pi)/2-log(varterm(i))/2-(dX(i)-dT(i)*mustep)^2/(2*varterm(i));
end

L=sum(Li);

f=-L;

end
```

**Table 2**

```
clear


mu_step=0.1;

var_step=0.1;

Vp=400;

y0=0;

Rerun=100;

M=1000

count_WLSbest=0;


for rerun=1:Rerun


for m=1:M

%% Generate data

y=y0*zeros(1,1000);

for t=2:1000

   tplot(t)=t;

   y(t)=y(t-1)+mu_step+sqrt(var_step)*randn;

end

tlabel=zeros(1,10);

ylabell=zeros(1,10);

err=zeros(1,10);


j=1;

for j=1:10

   for t=1:1000

      if t/100==j

         tlabel(j)=t-99*rand;

         j=j+1;

      end

   end

end

tlabel=round(tlabel);

for j=1:10

   n(1,j)=57*rand;

end
```

```
n=round(n)+3*ones(1,10);

j=1;
for t=1:1000
   if t==tlabel(j)
      individuals=sqrt(Vp)*randn(n(j),1);
      Mean(1,j)=mean(individuals);
      ytrue(j)=y(t);
      ylabell(j)=y(t)+Mean(1,j);
      err(j)=sqrt(Vp/n(j));
      if j==10 break
      end
      j=j+1;
   end
end

v=err.^2;
V=diag(v);

%% GRW

n_samples=ones(1,10);
var_samples=err.^2;

c=1;
Tsample=c*tlabel;

for i=1:9
   dT(i)=-Tsample(i)+Tsample(i+1);
   dX(i)=ylabell(i+1)-ylabell(i);
   nA(i)=n_samples(i);
   nD(i)=n_samples(i+1);
   varA(i)=var_samples(i);
   varD(i)=var_samples(i+1);
end

mustep_min=-100; mustep_max=100;
varstep_min=0; varstep_max=100;
```

```
par_lb=[mustep_min varstep_min];
par_ub=[mustep_max varstep_max];

% fmincon
par_guess=[0 0];
Aineq=[]; Bineq=[]; Aeq=[]; Beq=[];
fun_objective_handle=...
   @(par)fun_objective(par,dT,dX,varA,nA,varD,nD);
[par_opt,fval,exitflag,output,lambda,grad,hessian] =...
fmincon(fun_objective_handle,par_guess,Aineq,Bineq,Aeq,Beq,par_lb,par_ub);

mustep(m)=par_opt(1);
varstep(m)=par_opt(2);

b_GRW(m)=mustep(m);

X=ones(10,1);
a_GRW=inv(X'*inv(V)*X)*X'*inv(V)*(ylabell'-b_GRW(m)*tlabel');
for j=1:10
   yhat_GRW(j)=a_GRW+b_GRW(m)*tlabel(j);
end

WMSE_GRW(m)=(ylabell-yhat_GRW)*inv(V)*(ylabell-yhat_GRW)'/trace(inv(V));

%% WLS

X=[ones(10,1) tlabel'];
bls=inv(X'*inv(V)*X)*X'*inv(V)*ylabell';
a_WLS=bls(1);
b_WLS(m)=bls(2);
for i=1:10
   yhat_WLS(i)=a_WLS+b_WLS(m)*tlabel(i);
end

WMSE_WLS(m)=(ylabell-yhat_WLS)*inv(V)*(ylabell-yhat_WLS)'/trace(inv(V));

if varstep(m)==0
   b_WLS(m)=NaN;
```

```
    WMSE_WLS(m)=NaN;
end

if abs(varstep(m)-0.1)<0.01
% if abs(mustep(m)-0.1)<0.01
% if abs(mustep(m)-0.1)<0.01 & abs(varstep(m)-0.1)<0.01
    vartest(rerun)=varstep(m);
    b_GRWtest(rerun)=b_GRW(m);
    b_WLStest(rerun)=b_GRW(m);
    b_WLStest(rerun)=b_WLS(m);
    WMSE_GRWtest(rerun)=WMSE_GRW(m);
    WMSE_WLStest(rerun)=WMSE_WLS(m);
    if WMSE_WLStest(rerun)<WMSE_GRWtest(rerun);
        count_WLSbest=count_WLSbest+1;
    end
    break
end

end

end

Varstep=[mean(vartest) std(vartest)]
B_GRW=[mean(b_GRWtest) std(b_GRWtest)]
B_WLS=[mean(b_WLStest) std(b_WLStest)]

WMSE_GRW=[mean(WMSE_GRWtest) std(WMSE_GRWtest)]
WMSE_WLS=[mean(WMSE_WLStest) std(WMSE_WLStest)]

100*count_WLSbest/length(vartest)

%% Parameter search
function f = fun_objective(par,dT,dX,varA,nA,varD,nD)
mustep=par(1);
varstep=par(2);

for i=1:9
    varterm(i)=dT(i)*varstep+varA(i)/nA(i)+varD(i)/nD(i);
```

```
    Li(i)=-log(2*pi)/2-log(varterm(i))/2-(dX(i)-dT(i)*mustep)^2/(2*varterm(i));
end

L=sum(Li);

f=-L;

end
```

##### Table 3

```
clear

mu_step=0.1;
var_step=0.1;
Vp=400;
y0=0
countt=0
M=100

for m=1:M
%% Generate data
y=y0*zeros(1,1000);
for t=2:1000
    tplot(t)=t;
    y(t)=y(t-1)+mu_step+sqrt(var_step)*randn;
end
tlabel=zeros(1,10);
ylabell=zeros(1,10);
err=zeros(1,10);

j=1;
for j=1:10
    for t=1:1000
        if t/100==j
            tlabel(j)=t-99*rand;
            j=j+1;
        end
```

```
    end
end
tlabel=round(tlabel);
for j=1:10
    n(1,j)=57*rand;
end
n=round(n)+3*ones(1,10);

j=1;
for t=1:1000
    if t==tlabel(j)
        individuals=sqrt(Vp)*randn(n(j),1);
        Mean(1,j)=mean(individuals);
        ytrue(j)=y(t);
        ylabell(j)=y(t)+Mean(1,j);
        err(j)=sqrt(Vp/n(j));
        if j==10 break
        end
        j=j+1;
    end
end

v=err.^2;
V=diag(v);

%% GRW

n_samples=ones(1,10);
var_samples=v;

c=1;
Tsample=c*tlabel;

for i=1:9
    dT(i)=Tsample(i+1)-Tsample(i);
    dX(i)=ylabell(i+1)-ylabell(i);
    nA(i)=n_samples(i);
    nD(i)=n_samples(i+1);
```

```
   varA(i)=var_samples(i);
   varD(i)=var_samples(i+1);
end

mustep_min=-100; mustep_max=100;
varstep_min=0; varstep_max=100;
par_lb=[mustep_min varstep_min];
par_ub=[mustep_max varstep_max];

% fmincon
par_guess=[0 0];
Aineq=[]; Bineq=[]; Aeq=[]; Beq=[];
fun_objective_handle=...
   @(par)fun_objective(par,dT,dX,varA,nA,varD,nD);
[par_opt,fval,exitflag,output,lambda,grad,hessian] =...
fmincon(fun_objective_handle,par_guess,Aineq,Bineq,Aeq,Beq,par_lb,par_ub);

mustep(m)=par_opt(1);
varstep(m)=par_opt(2);

if varstep(m)<0.00001
   countt=countt+1;
end

for i=1:9
   varterm(i)=dT(i)*varstep(m)+varA(i)/nA(i)+varD(i)/nD(i);
   Li(i)=-log(2*pi)/2-log(varterm(i))/2-(dX(i)-dT(i)*mustep(m))^2/(2*varterm(i));
end

L=sum(Li);

k=2;
N=9;

AICc_GRW(m)=2*k-2*L+2*k*(k+1)/(N-k-1);

X=ones(10,1);
b_GRW(m)=mustep(m);
```

```
a_GRW(m)=inv(X'*inv(V)*X)*X'*inv(V)*(ylabell'-b_GRW(m)*tlabel');
for j=1:10
   yhat_GRW(j)=a_GRW(m)+b_GRW(m)*tlabel(j);
end

WMSE_GRW(m)=((ylabell-yhat_GRW)*inv(V)*(ylabell-yhat_GRW)')/trace(inv(V));

%% WLS

X=[ones(10,1) tlabel'];
bls=inv(X'*inv(V)*X)*X'*inv(V)*ylabell';
a_WLS(m)=bls(1);
b_WLS(m)=bls(2);
for i=1:10
   yhat_WLS(i)=a_WLS(m)+b_WLS(m)*tlabel(i);
end

WMSE_WLS(m)=(ylabell-yhat_WLS)*inv(V)*(ylabell-yhat_WLS)'/(trace(inv(V)));

for j=1:10
  lnw(j)=log(sqrt(v(j)));
  z(j)=(ylabell(j)-yhat_WLS(j)).^2/v(j);
end

k=3;
N=10;

AICc_WLS(m)=2*k+N*log(2*pi)+N+2*sum(lnw)+N*log(sum(z)/N)+2*k*(k+1)/(N-k-1);

end

Varstep=[mean(varstep) std(varstep)]
BGRW=[mean(b_GRW) std(b_GRW)]
BWLS=[mean(b_WLS) std(b_WLS)]

AICcGRW=[mean(AICc_GRW) std(AICc_GRW)]
AICcWLS=[mean(AICc_WLS) std(AICc_WLS)]
```

```
[varstep' AICc_GRW'-AICc_WLS' WMSE_GRW'-WMSE_WLS']

100*countt/M

%% Parameter search
function f = fun_objective(par,dT,dX,varA,nA,varD,nD)
mustep=par(1);
varstep=par(2);

for i=1:9
   varterm(i)=dT(i)*varstep+varA(i)/nA(i)+varD(i)/nD(i);
   Li(i)=-log(2*pi)/2-log(varterm(i))/2-(dX(i)-dT(i)*mustep)^2/(2*varterm(i));
end

L=sum(Li);

f=-L;

end
```
